# CRISPR-enhanced human adipocyte “browning” as cell therapy for metabolic disease

**DOI:** 10.1101/2020.10.13.337923

**Authors:** Emmanouela Tsagkaraki, Sarah Nicoloro, Tiffany De Souza, Javier Solivan-Rivera, Anand Desai, Yuefei Shen, Mark Kelly, Adilson Guilherme, Felipe Henriques, Raed Ibraheim, Nadia Amrani, Kevin Luk, Stacy Maitland, Randall H. Friedline, Lauren Tauer, Xiaodi Hu, Jason K. Kim, Scot A. Wolfe, Erik J. Sontheimer, Silvia Corvera, Michael P. Czech

## Abstract

Obesity and type 2 diabetes (T2D) are associated with poor tissue responses to insulin^1,2^, disturbances in glucose and lipid fluxes^3–5^ and comorbidities including steatohepatitis^6^ and cardiovascular disease^7,8^. Despite extensive efforts at prevention and treatment^9,10^, diabetes afflicts over 400 million people worldwide^11^. Whole body metabolism is regulated by adipose tissue depots^12–14^, which include both lipid-storing white adipocytes and less abundant “brown” and “brite/beige” adipocytes that express thermogenic uncoupling protein UCP1 and secrete factors favorable to metabolic health^15–18^. Application of clustered regularly interspaced short palindromic repeats (CRISPR) gene editing^19,20^ to enhance “browning” of white adipose tissue is an attractive therapeutic approach to T2D. However, the problems of cell-selective delivery, immunogenicity of CRISPR reagents and long term stability of the modified adipocytes are formidable. To overcome these issues, we developed methods that deliver complexes of SpyCas9 protein and sgRNA *ex vivo* to disrupt the thermogenesis suppressor gene *NRIP1*^21,22^ with near 100% efficiency in human or mouse adipocytes. *NRIP1* gene disruption at discrete loci strongly ablated NRIP1 protein and upregulated expression of UCP1 and beneficial secreted factors, while residual Cas9 protein and sgRNA were rapidly degraded. Implantation of the CRISPR-enhanced human or mouse brown-like adipocytes into high fat diet fed mice decreased adiposity and liver triglycerides while enhancing glucose tolerance compared to mice implanted with unmodified adipocytes. These findings advance a therapeutic strategy to improve metabolic homeostasis through CRISPR-based genetic modification of human adipocytes without exposure of the recipient to immunogenic Cas9 or delivery vectors.

---

The use of human cells as therapeutics offers major advantages over small molecule drugs and biologics in treating certain diseases based on their abilities to home to specific organs or cell types, initiate cell-cell interactions and secrete multiple bioactive factors^23,24^. Although still in early stages of development, cellular therapies have already had major impact on treatment of certain forms of cancer such as leukemia, lymphoma, melanoma and small cell lung carcinoma^25,26^. This approach involves genetic modification *ex vivo* of immune cells taken from a human subject to enhance their ability to disrupt malignancies upon infusion back into the same subject. In theory, this strategy should be effective in diseases in which cells with relevant therapeutic potential can be genetically modified to enhance that potential. Here we take advantage of recent discoveries revealing the utility of thermogenic adipocytes to function as major beneficial regulators of whole body metabolism in such metabolic diseases as type 2 diabetes and obesity^15–18^. Thermogenic adipocytes, denoted as brown^27^, beige^28^ or brite^27,29^, are distinct from the more abundant lipid storing white adipocytes not only by their high oxidative capacity and expression of mitochondrial uncoupling protein (UCP1) but also by their secretion of factors that enhance energy metabolism and energy expenditure^15–18^. Multiple studies have demonstrated that implantation of mouse brown adipose tissue into obese, glucose intolerant mice can improve glucose tolerance and insulin sensitivity^30–32^. Recently, human beige adipocytes expanded from small samples of subcutaneous adipose tissue were shown to form robust thermogenic adipose tissue depots upon implantation into immune-compromised obese mice and to lower blood glucose levels^33^. Collectively, these data provide the framework to apply genetic modifications to adipocytes to further improve their therapeutic potential.

## SpyCas9/sgRNA RNPs for *ex vivo* gene editing

In order to enhance the therapeutic potential of adipocytes in obesity and diabetes, we initially targeted the mouse *Nrip1* gene. *Nrip1* had been previously shown to strongly suppress glucose transport, fatty acid oxidation, mitochondrial respiration, uncoupling protein 1 (UCP1) expression as well as secretion of such metabolically beneficial factors including neuroregulin 4^21,22,34^. NRIP1 functions as a transcriptional co-repressor that attenuates activity of multiple nuclear receptors involved in energy metabolism, including estrogen related receptor (ERRα), peroxisome proliferator activated receptor (PPARγ) and thyroid hormone receptor (TH)^35^. NRIP1 knockout in white adipocytes upregulates genes that are highly expressed in brown adipocytes, enhancing glucose and fatty acid utilization and generating heat. *Nrip1* ablation in mice elicits a lean phenotype under high fat diet conditions, and greatly enhances energy expenditure, glucose tolerance and insulin sensitivity^21^. However, NRIP1 is not an attractive target for conventional pharmacological intervention as it is not an enzyme and has a multiplicity of tissue specific roles such as regulating the estrogen receptor in the reproductive tract^35^. Targeting NRIP1 selectively within adipocytes represents an ideal approach to capture its therapeutic potential without undesirable side effects.

A key aspect of our strategy in targeting the *Nrip1* gene was to employ methods that would ablate its expression in adipocytes but not cause immune responses upon implantation of the cells. CRISPR-based methods based on continuous expression of Cas9/sgRNA to modify adipocytes that function *in vivo* have been reported^36,37^, but they expose recipients to Cas9 and delivery agents that cause immune responses. Direct administration of Cas9/sgRNA complexes in mice have not been adipocyte-specific and could cause undesirable effects in other tissues^36^. Ribonucleoprotein complexes of SpyCas9/sgRNA are desirable vehicles for such modifications since they are rapidly degraded following DNA disruption^38^. A previous attempt at delivery of such CRISPR-based complexes to adipocytes were suboptimal as efficiencies of delivery of RNPs to these cells was only modest^34^. We overcame these deficiencies by disrupting *Nrip1* in mouse preadipocytes with ribonucleoprotein (RNP) complexes of Cas9 and sgRNA by modifying electroporation methods^39^ described for other cell types (Extended Data Fig 1), and confirmed Cas9 protein is rapidly degraded following indel formation in preadipocytes(Extended Data Fig. 2). Electroporation conditions were developed to optimize the efficiency of *Nrip1* gene targeting in mouse preadipocytes by Cas9/sgRNA RNPs without perturbing their differentiation into adipocytes (Fig. 1a,b and Extended Data Fig. 1).

**Figure 1.**
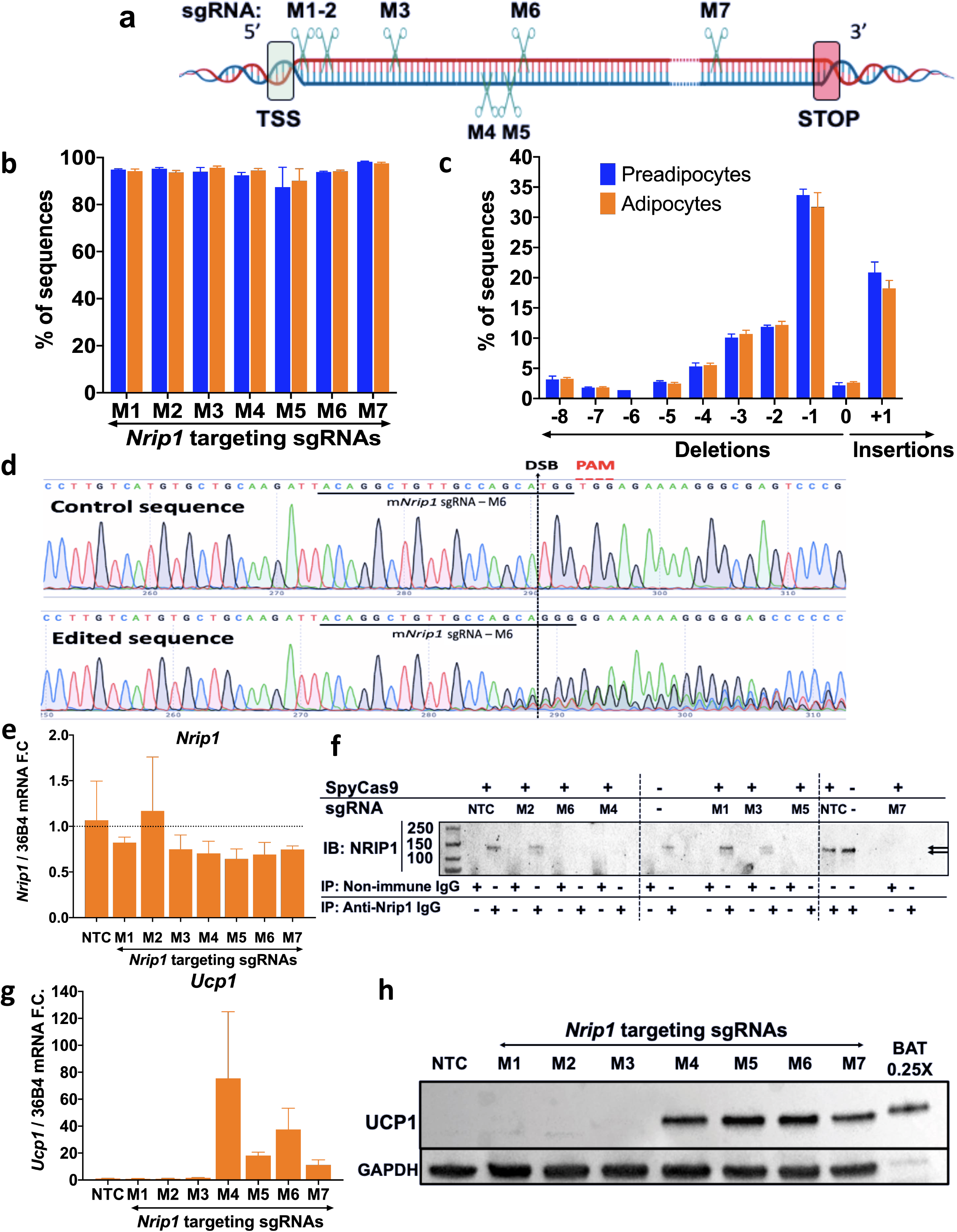
High efficiency *Nrip1* gene disruption at 7 loci by SpyCas9/sgRNA RNPs produces variable degrees of NRIP1 protein loss and UCP1 upregulation in murine primary adipocytes. **a.** Mapping of the sgRNAs M1-M7 targeting various loci of murine *Nrip1* coding region which is entirely located in exon 4 (TSS= transcription start site; STOP = stop codon). **b.** Editing efficiency as evaluated with indel percentage 72 hours after the transfection of primary preadipocytes (blue) and differentiation to mature primary adipocytes (orange). **c**. Indel distribution of *Nrip1* sgRNA-M6 with frameshift indels that are sustained after differentiation. **d**. Sanger sequencing traces of control vs *Nrip1* disrupted cells with sgRNA-M6 showing the sgRNA binding site (solid black line), PAM (red), the double-strand-break (denoted as DSB) site (dashed black line) on the sgRNA-M6 targeting locus and the traces downstream of the DSB created by the DNA repair mechanisms (figure created with SnapGene). **e**. *Nrip1* gene expression detected by RT-PCR in mature adipocytes targeted with the different sgRNAs. **f**. Immunoprecipitation assay for NRIP1 (140 kDa, arrows at right) in mature primary adipocytes on day 6 post differentiation targeted with the different sgRNAs. The total lysate protein amount used in the assay was 250 μg per sample. **g**. *Ucp1* expression by RT-PCR in mature adipocytes targeted with the different sgRNAs compared to non-targeted control cells. **h**. Western blot for UCP1 protein (33kDa) in mature adipocytes on day 6 post differentiation targeted with the different sgRNAs. Lanes 1-8 were loaded with 20 μg of total protein while lane 9 was loaded with 5 μg οf total protein isolated from mouse BAT. NTC = Non-targeting control. In panels b and c error bars denote Mean ± S.E.M. In panels e and g, bars denote mean, error bars denote mean ± standard deviations, n ≥ 3 biological replicates.

The efficiencies of indel formation by 7 different sgRNAs against various regions of *Nrip1* gene exon 4 (Fig. 1a) were uniformly sustained in the 90% range in preadipocytes and upon their differentiation into adipocytes (Fig. 1b,c). Indels were quantified by Sanger sequencing data analysis of PCR fragments spanning the upstream and downstream double stranded breaks of the *Nrip1* genomic DNA (Fig. 1c,d) with little change in the total Nrip1 mRNAs(Fig 1e). High frequencies of frameshift mutations in *Nrip1* by all 7 sgRNAs were found and similar indels were found in the corresponding *Nrip1* mRNA species, as exemplified by sgRNA M3 and M4 (Extended Data Fig. 3).

While the mRNA of *Nrip1* was equally abundant in all groups, indicating no increased degradation due to disruption (Fig. 1e), surprisingly, not all of the sgRNAs were effective in eliciting loss of the NRIP1 protein (Fig. 1f). Consistent with these data, thermogenic responses to the various sgRNAs as reflected by elevated expression of UCP1 mRNA (Fig. 1g) and protein (Fig. 1h) correlated with the loss of native full length NRIP1 protein. Taken together, these data show that sgRNAs targeting the regions of *Nrip1* DNA that encode the N-terminal region of the NRIP1 protein are not effective in eliminating synthesis of functional NRIP1 protein. Most likely, additional transcription or translation start sites beyond these target sites are functional under these conditions. Thus, sgRNAs that are optimal for inducing thermogenic genes must be identified by such screening methods.

## Implantation of CRISPR-modfied mouse adipocytes

To test the ability of NRIP1-deficient adipocytes to improve metabolism in mice, large numbers of primary preadipocytes obtained from 2-3 week old mice were electroporated with RNPs consisting of either SpyCas9/non targeting control (NTC) sgRNA or SpyCas9/sgRNA-M6 complexes, and then differentiated into adipocytes and implantated into wild type mice. The implanted mice were kept on normal diet for 6 weeks during the development of adipose tissue depots from the injected adipocytes, then placed on a high fat diet (HFD) regimen to enhance weight gain (Fig. 2a). Adipocytes treated with Cas9/sgRNA-M6 displayed upregulation of *Ucp1* and other genes highly expressed in brown adipocytes (e.g., *Cidea*) prior to transplantation (Extended Data Fig. 4). A transient decrease in overall body weights were detected between mice implanted with RNPs containing the Cas9/sgRNA-M6 versus the Cas9/NTCsgRNA group, but by 6 weeks of HFD no significant difference was observed (Fig. 2b). Nonetheless, implantation of NRIP1KO adipocytes prevented the increase in fasting blood glucose concentration due to HFD that occurs in the control adipocyte-implanted mice (Fig. 2c). Glucose tolerance was also significantly improved by implantation of NRIP1KO adipocytes (Fig 2d,e). The implanted adipose tissue depots retained their elevated expression of UCP1 16 weeks after implantation, at which time they were excised for analysis (Fig. 2f). The livers and inguinal white adipose tissues (iWAT) from the Cas9/sgRNA-M6 group of mice had lower weights (Fig. 2g) and lower iWAT to body weight ratios (Extended Data Fig. 4), revealing a strong systemic effect.

**Figure 2.**
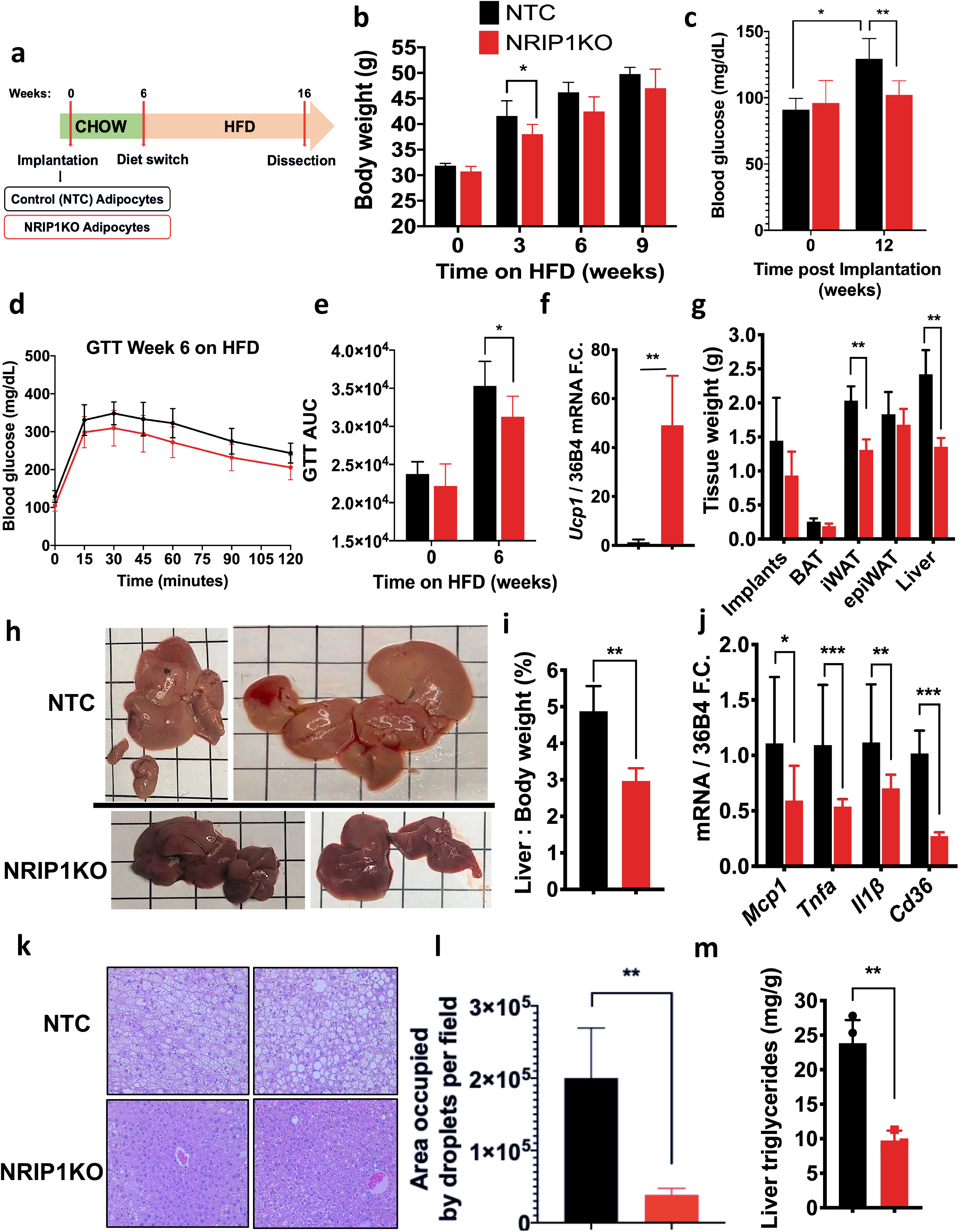
Implantation of NRIP1-depleted mouse adipocytes improves glucose tolerance and markedly decreases iWAT weight and liver triglyceride accumulation in recipient mice. Mice were implanted with either mouse adipocytes previously treated with NTC sgRNA/Cas9 RNPs or with sgRNA-M6/Cas9 RNPs (NRIP1KO). **a**. Schematic protocol of implantation of murine NTC adipocytes or NRIP1KO adipocytes into C57BL/6 wild type mice followed by 60% kCal fat diet (HFD). **b**. Total body weights of recipients on chow after implantation and after 3, 6 and 9 weeks on HFD. **c.** Fasting blood glucose concentrations at 12 weeks post implantation (6 weeks after start of HFD). **d**. Glucose tolerance test *(GTT) after 16-hour overnight fasting in implant recipients after 6 weeks on HFD. **e.** Bar graphs of areas under the curve from GTTs in implant recipient mice on chow and after 6 weeks on HFD. **f.** *Ucp1* expression in the implants harvested after the dissection at study termination. **g**. Weight of total bilateral implants, whole BAT, total bilateral iWAT, epiWAT and total liver as measured after dissection. **h**. Macroscopic images of the whole livers of the implant recipients after dissection (square = 1cm^2^). **i**. Liver over whole body weight percentage. **j**. Expression of genes related to inflammation and involved in hepatic steatosis in the livers of implant recipients detected by RT-PCR. **k**. Hematoxylin and eosin (H&E) stain on liver histology of the implant recipients at 20X magnification. **l**. Quantification of total H&E images of implant recipients’ livers for total area occupied by lipid droplets per field. **m**. Triglyceride measurements in pulverized liver extracts after dissection. Panel c-e: black = NTC adipocytes; red = NRIP1KO adipocytes, n ≥ 3 biological replicates, panels c-e: black = NTC cell implants (n=7); red = NRIP1KO cell implants (n=8), Panels b,f-m: black = NTC cell implant recipients (n=4); red = NRIP1KO cell recipients (n=3). Bars represent the mean, error bars denote mean ± standard deviation. * p < 0.05, ** p < 0.01, *** p < 0.001 by unpaired two-tailed T-test.

Livers of the NRIP1 deficient adipocyte-implanted mice were less pale (Fig. 2h), were smaller as assessed by lower liver to body weight ratios (Fig. 2i) and displayed lower expression of genes associated with fat metabolism (CD36) and with inflammation (Mcp1, Tnfα, II1β) (Fig. 2j) compared to mice implanted with control adipocytes. Lipid droplets in the livers of mice with implants of Cas9/sgRNA-M6-treated adiocytes (Fig. 2k) were greatly decreased as assessed by quantification of lipid droplet area (Fig. 2l), size and number (Extended Data Fig. 4) as well as liver triglyceride determination (Fig. 2m). The decrease in hepatic lipid accumulation and inflammation in response to implantation of the NRIP1 depleted adipocytes suggests that this therapeutic approach might mitigate these T2D co-morbidities in humans^6^.

## Translation to human adipocytes

To translate these CRISPR-based methods to human adipocytes, adipocyte progenitors were obtrained from small samples of excised subcutaneous adipose tissue as previously described^33^. Electroporation conditions were tested to optimize efficiency of indel formation using various sgRNAs directed against regions of the *NRIP1* exon 4 at locations roughly similar to those we targeted in the mouse genomic DNA (compare Fig. 1a to Fig. 3a). Efficiencies of *NRIP1* gene disruption were observed with several sgRNAs in the 90% range (Fig. 3b), with indel distributions very similar in preadipocytes and adipocytes (Fig. 3c). Electroporated, NRIP1 deficient human preadipocytes could be readily differentiated to adipocytes without apparent disruption following indel formation (Fig. 3d). *NRIP1* mRNA was equally abundant in all conditions (Fig. 3e). *UCP1* expression increased by up to 100 fold in several experiments in Cas9/sgRNA treated cells, but only with certain sgRNAs (Fig. 3f,g), similar to our results with mouse adipocytes. Functional NRIP1 protein can still be expressed despite high efficiency indel formation in the N-terminal region. *NRIP1* disruption in combination with the adenylate cyclase activator forskolin synergistically increased UCP1 expression (Fig. 3h).

**Figure 3.**
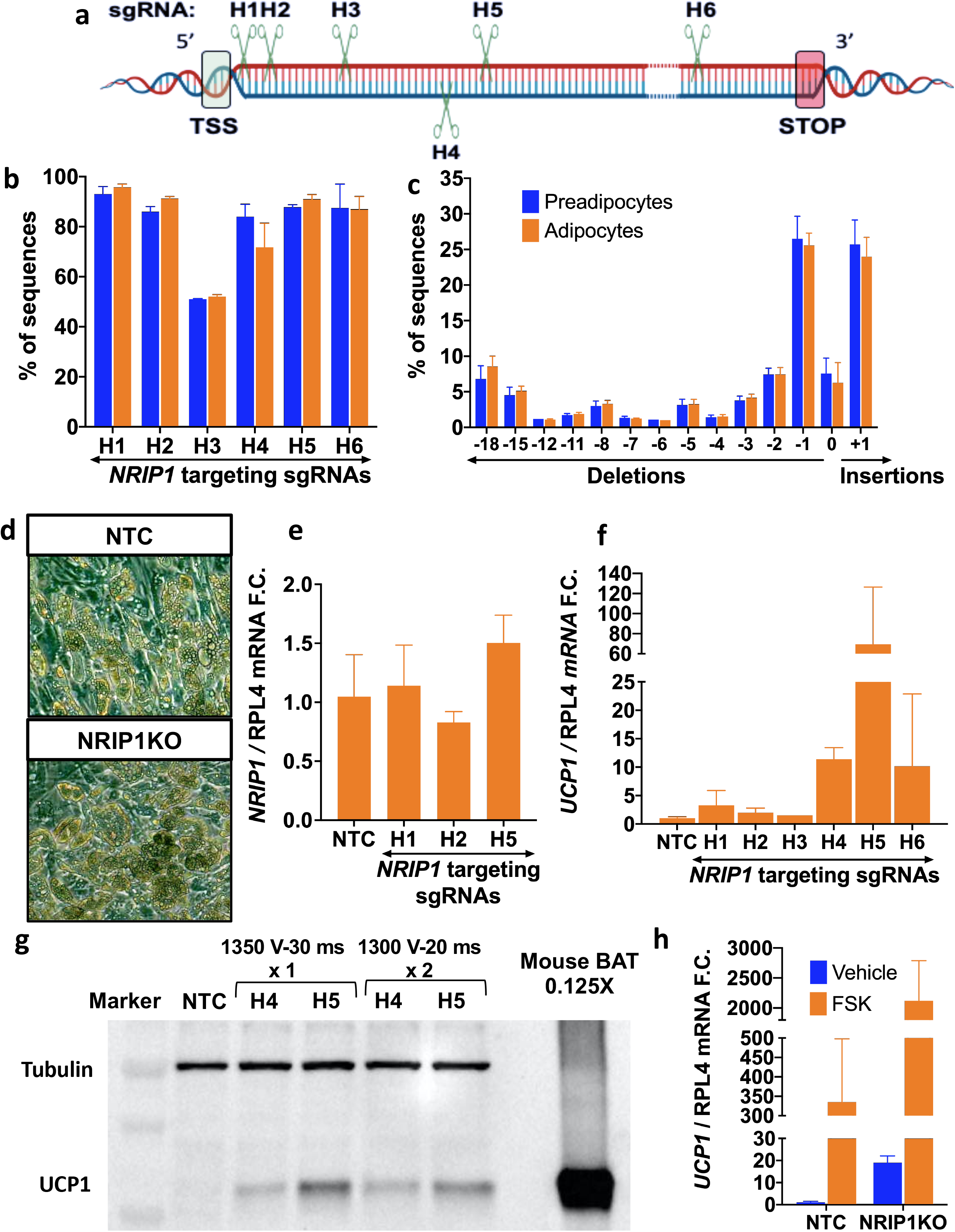
High efficiency *NRIP1* disruption in human adipocytes by SpyCas9 reveals variable UCP1 upregulation in a screen of sgRNAs targeting different loci of *NRIP1*. **a**. Mapping of sgRNAs H1-H6 screened against human *NRIP1* coding region entirely located in exon 4. **b**. Editing efficiency as evaluated with indel percentage 72 hours after the transfection of primary preadipocytes (blue) and differentiation to mature primary adipocytes (orange). **c**. Indel distribution of *NRIP1* sgRNA-H5 with frameshift indels that are sustained after differentiation. **d**. Microscopic image of cell culture of non-targeted control (top) and *NRIP1* disrupted (bottom) mature adipocyte morphology in cell culture at 10X magnification. **e**. *NRIP1* gene expression by RT-PCR in mature adipocytes on day 7 post differentiation targeted with the different sgRNAs. **f**. *UCP1* expression by RT-PCR in mature adipocytes on day 7 post differentiation targeted with the different sgRNAs compared to non-targeted control cells. **g**. Western blot for UCP1 protein (33kDa) in mature adipocytes on day 7 post differentiation targeted by the sgRNAs H4 and H5 with two different electroporation optimization protocols. Lanes 2-6 were loaded with 20 μg of total protein while lane 9 was loaded with 2.5 μg of total protein isolated from mouse BAT. **h**. *UCP1* gene expression by RT-PCR in non-targeted control or NRIP1 depleted adipocytes on day 7 post differentiation after a 7-hour stimulation of forskolin 10 μM or vehicle. NTC = Non-targeting control. Bars denote mean. In panels b and c, error bars denote Mean ± S.E.M. In panels e,f and h error bars denote mean ± standard deviations, n ≥ 3 biological replicates.

To test the efficacy of NRIP1-depleted human adipocytes to provide metabolic benefits in obese glucose intolerant mice, we utilized immune-compromised NOD.Cg-Prkdc^scid^ II2rg^tm1Wjl^/SzJ (NSG) mice that lack T cells, B cells and natural killer cells to accept human cell implants in the protocol depicted (Fig. 4a,b). *NRIP1* gene disruption was about 80% with the sgRNA-H5 (Fig 4c) and circulating human adiponectin (Fig. 4d) was the same from NTC vs sgRNA-H5 groups, indicating similar levels of adipose tissue formation. NRIP1KO adipocytes exhibited upregulation of UCP1 levels similar to Fig. 3 (not illustrated). and the resulting NRIP1KO adipose tissue implants harvested from the mice 13 weeks later retained the enhanced UCP1 expression (not shown).

While no body weight difference between groups was detected on normal diet, a highly significant decrease in weight gain on the HFD was observed in mice implanted with *NRIP1* disrupted adipocytes (Fig. 4e). Importantly, mice with control human adipocyte implants displayed significantly decreased glucose tolerance 3 weeks after starting a HFD while animals with NRIP1-depleted adipocyte implants did not (Fig. 4f-i). The difference in glucose tolerance between the two groups at the end of the study was highly significant(Fig. 4h,i). Relative liver to body weight ratios (Fig 4j) and liver triglycerides (Fig. 4k) were also decreased when implants were performed with NRIP1 deficient adipocytes compared to control adipocytes.

**Figure 4.**
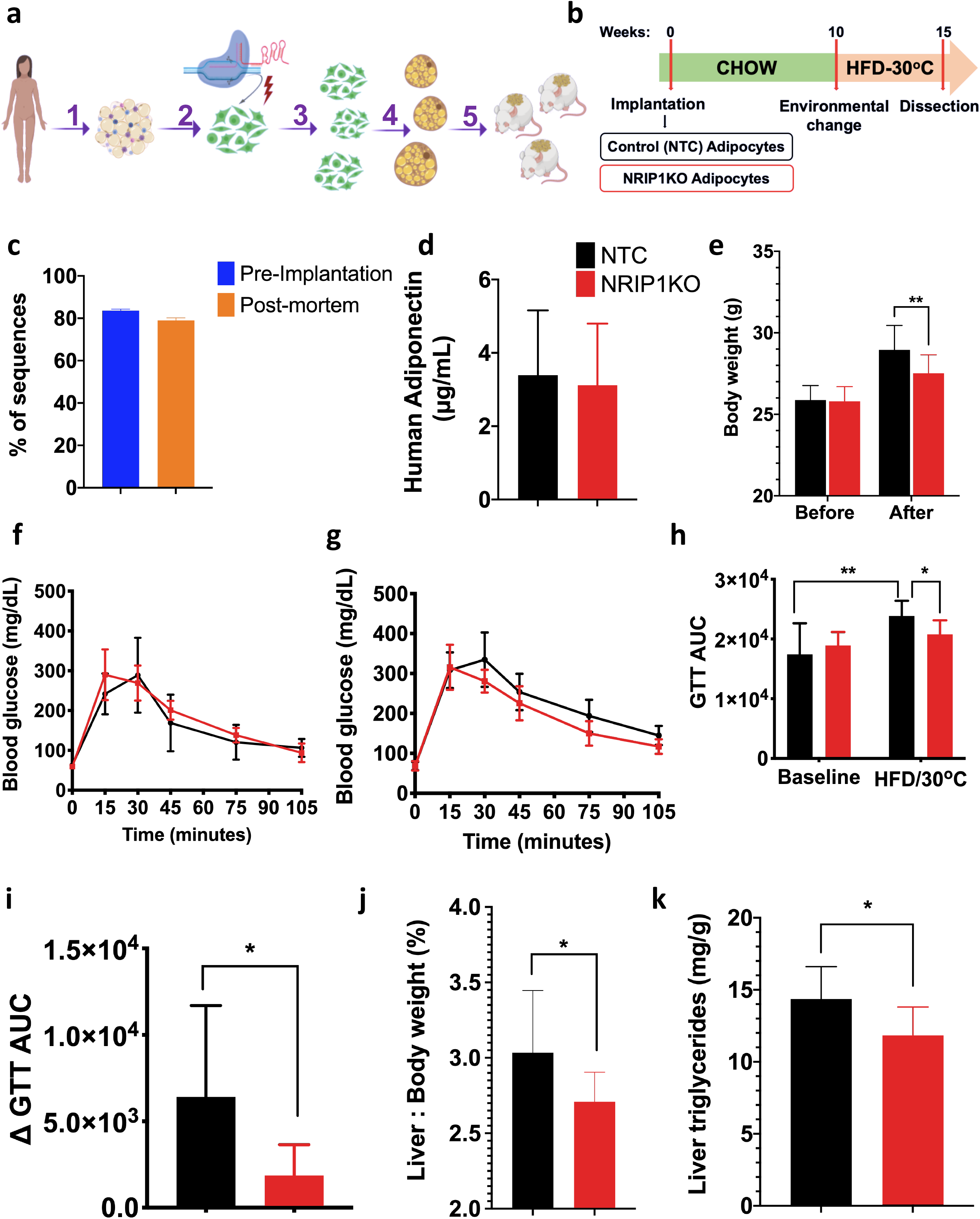
Implantation of NRIP1-targeted human adipocytes decreases body weight as well as liver triglyceride, and enhances glucose tolerance in recipient immunocompromised, HFD fed NSG mice. Mice were implanted with either human adipocytes previously treated with NTC sgRNA/Cas9 RNPs or with sgRNA-H5/Cas9 RNPs (NRIP1KO). **a**. Study description: 1) adipose tissue isolation from a human donor during panniculectomy, 2) harvesting of human primary preadipocytes after development Cas9/sgRNA RNPs were delivered into the human preadipocytes by electroporation followed by 3) expansion 1:6 of the genetically modified preadipocytes and 4) their differentiation into mature adipocytes; 5) Implantation of non-targeted control (NTC) sgRNA treated adipocytes versus the NRIP1 depleted adipocytes was performed in the dorsal area of NSG mice. **b**. Schematic protocol of implantation of human NTC adipocytes or NRIP1KO adipocytes into NSG mice followed by HFD feeding. **c**. Editing efficiency as evaluated with percentage of indels in the mature adipocytes transfected with NRIP1KO sgRNA-H5 before implantation (blue) and indel percentage in the genomic DNA isolated from the NRIP1 depleted implants 15 weeks following transplantation. **d**. Human adiponectin levels detected in the plasma of NSG recipients 9 weeks following transplantation for the assessment of engraftment and functionality of the implants. **e**. Total body weight of NSG mouse recipients before transplantation (left) and after 3 weeks on HFD and thermoneutrality (right). **f**. Baseline glucose tolerance test after 16 hr fasting before transplantation. **g**. Glucose tolerance test after 3 weeks on HFD and thermoneutrality. **h**. Glucose tolerance test areas under the curve (GTT AUC) before transplantations (left) and 3 weeks after HFD under thermoneutrality (right). **i**. Matched difference of the GTT AUC before transplantations (left) and 3 weeks after HFD and thermoneutrality (right). **j**. Liver over whole body weight percentage. **k**. Triglyceride measurements in pulverized liver extracts after dissection. Black = NTC cell implant recipients (n=4); red = NRIP1KO cell implant recipients (n=6) Bars denote mean, error bars denote mean ± standard deviation. * p < 0.05, ** p < 0.01 by unpaired T-test.

Taken together, the data presented here show that CRISPR-based RNPs can enhance “browning” of adipocytes *ex vivo* at high efficiency without the use of expression vectors to improve metabolic parameters in two mouse models of obesity. Although CRISPR-based upregulation of UCP1 alone in implanted adipocytes can improve metabolism in mice^37^, targeting *NRIP1* has the advantage of upregulating expression of many genes that have favorable metabolic effects in addition to UCP1. The specific approach presented here has several additional advantages, including the rapid electroporation procedure with minimal loss of viability, the fact that NRIP1 deletion does not diminish adipose differentiation, and the lack of immunogenic Cas9/sgRNA complexes in implanted cells. Also, the brief exposure of cells to Cas9/sgRNA that we document here (Extended Fig. 2) reduces the potential for off target effects that are produced by long term expression of these reagents. Since UCP1 expression in NRIP1KO adipocytes does not reach the level of mouse BAT, our approach can be improved by further enhancing adipocyte browning through disrupting combinations of targets in addition to NRIP1. Indeed we find that simultaneous delivery of multiple sgRNAs into preadipocytes each yield high efficiencies of indel formation (not shown), offering the potential for disrupting multiple thermogenic suppressor genes to achieve greater therapeutic potential. Also the use of high fidelity nucleases^40,41^ can further maximize cell viability and functionality. Further experiments to define the dose of implanted cells and preferential recipient sites will also help optimize this technique as a step towards testing in larger animals and advancement towards clinical trials.

## Acknowledgememts

We wish to thank members of the Czech and Corvera laboratories for helpful discussions and Kerri Miller for excellent assistance in preparing the manuscript. We thank the University of Massachusetts Morphology Core Facility for assistance in the histological preparations, stains and analysis. This work was supported by the Assistant Secretary of Defense for Health Affairs endorsed by the Department of Defense, through the Peer Reviewed Medical Research Program under Award No. W81XWH-18-1-0397 and W81XWH-18-1-0398 (to S.C. and M.P.C.). Opinions, interpretations, conclusions and recommendations are those of the authors and are not necessarily endorsed by the Department of Defense.This work was also supported by National Institutes of Health grants DK030898 (to M.P.C.), GM115911 (to E.J.S. and S.A.W.), TR002668 (to E.J.S. and S.A.W.), HL147482 (to K.L.), the the UMASS Mouse Metabolic Phenotyping Center (NIH grant 5U2C-DK093000). We also gratefully acknowledge generous funding through the Isadore and Fannie Foxman Chair in Medical Science (to M.P.C.), the Endowed Professorship in Diabetes Research Chair (to S.C.) and postdoctoral fellowship support to Felipe Henriques by the American Diabetes Foundation (Grant 1-19-PMF-035). Some figures were created with Biorender.com.

## Author Contributions

E.T., S.N., S.C. and M.P.C. designed the study and wrote the manuscript. E.T. and S.N. performed most of the experiments, analyzed the data and performed the mouse adipocyte implantation studies. E.T., T.D.S., J.SR. performed the human adipocyte implantation studies. T.D.S., J.SR. and A.D. established the human adipose explant-derived cell lines. Y.S. contributed to the initial strategy of the work. M.K. established the colonies for mouse cell donors and performed the blood collections of the recipient mice. A.G. and F.H. performed experiments and guided methods. E.T., R.R.I., N.A. and E.J.S. contributed to the gene-editing optimization and cloning experiments and to interpretation of data. K.L., S.M. and S.A.W. purified SpyCas9 protein and developed the plasmids. R.H.F., L.T., X.H. and J.K.K. contributed to strategies of evaluating metabolic parameters. All authors reviewed and were invited to edit the manuscript.

## Extended data

**Extended Data Figure 1.**
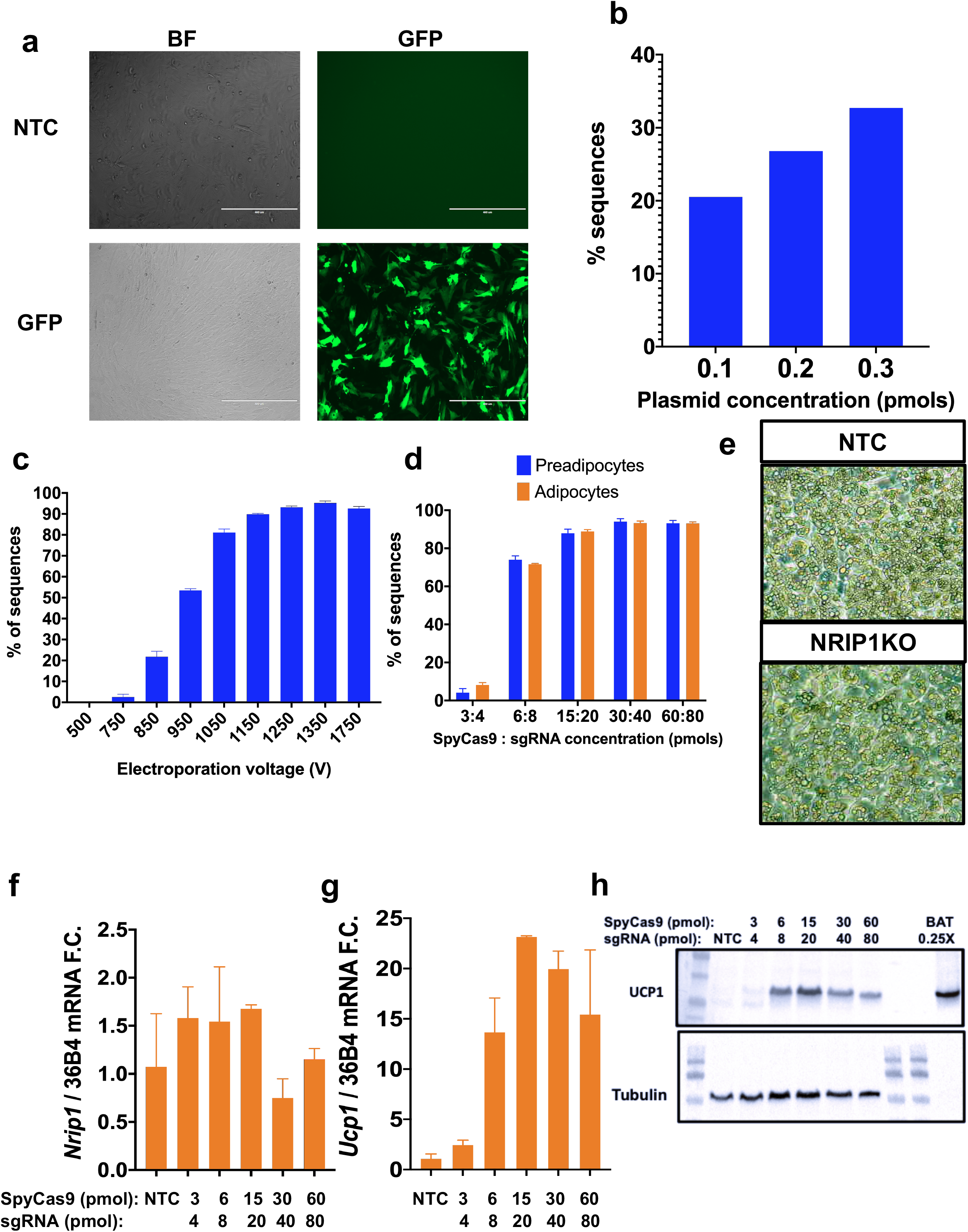
SpyCas9/sgRNA RNPs are more efficiently delivered into murine adipocytes by electroporation than plasmid expression. Delivery of SpyCas9/sgRNA RNPs versus plasmids encoding either GFP or Cas9 and sgRNA electroporated with preadipocytes were compared. **a**. GFP expression plasmid (Lonza pmaxGFP) was used in various concentrations for optimization of the plasmid delivery by electroporation in murine preadipocytes (data not shown) to determine the optimum range (0.1-0.2 pmols) that induces GFP expression by fluorescent microscopy (white line = 400 μm). Top: Control preadipocytes Bottom: Preadipocytes transfected with 0.2 pmols at 72 hours after electroporation (1350 V, 30 ms, 1 pulse). **b.** Application of the optimized electroporation protocol to transfect preadipocytes with plasmids expressing SpyCas9 and sgRNA-M6 in three different concentrations within the range determined with the GFP plasmid titration: 0.1 - 0.3 pmols. **c.** Optimization of the electroporation protocol to deliver CRISPR RNPs in preadipocytes with 1 pulse and 30ms width and increasing voltage. **d.** Titration of RNP concentrations in correlation with the editing efficiency with sgRNA-M6. **e.** Mature adipocytes transfected before differentiation with either NTC RNPs or sgRNA-M6 in cell culture and magnification 10X. **f.** *Nrip1* gene expression by RT-PCR in mature adipocytes transfected with varying RNP concentrations. **g.** *Ucp1* gene expression by RT-PCR in mature adipocytes transfected with varying RNP concentrations. **h.** UCP1 expression by western blot in mature adipocytes transfected with various RNP concentrations. NTC = Non-targeting control. Bars denote mean. In figures c, d error bars denote Mean ± S.E.M. In panels f, g error bars denote mean ± standard deviations. n ≥ 3 biological replicates. Nrip1 sgRNA-M5 was used in the titration experiment.

**Extended Data Figure 2.**
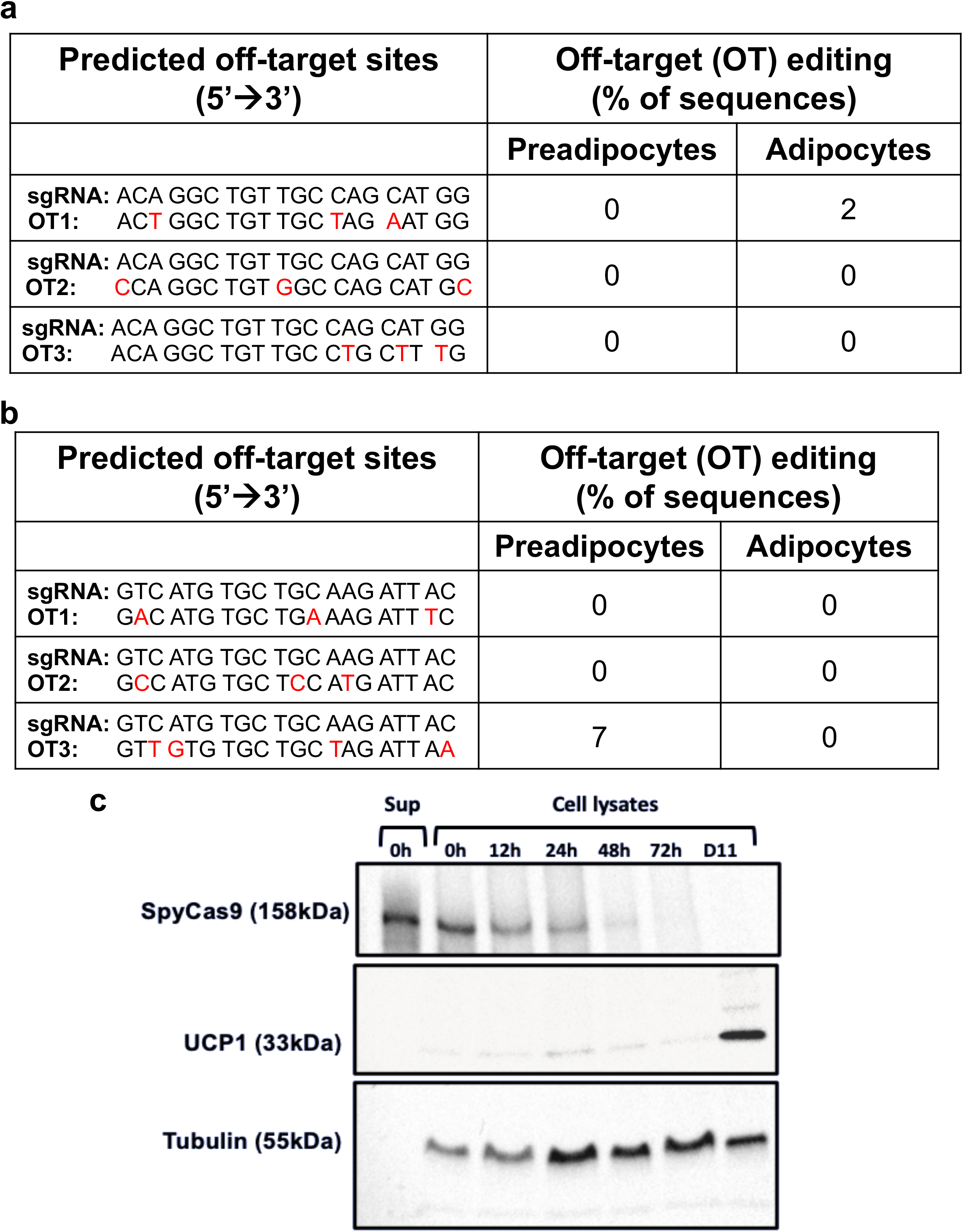
SpyCas9/sgRNA RNP-mediated *Nrip1* disruption in preadipocytes is characterized by little or no off-target editing and degradation of SpyCas9. **a.** Editing at the top three predicted off-target loci of the sgRNA-M6 in the murine genome by sanger sequencing data analysis. **b.** Editing at the top three predicted off-target loci of the sgRNA-H5 in the human genome by sanger sequencing data analysis. **c.** Time-course SpyCas9 protein detection by Western Blotting in cell lysates at various time points after electroporation to deliver RNPs of SpyCas9: sgRNA 30:40 pmols (sup 0 hours), in preadipocytes immediately after electroporation (0 hours), in preadipocytes at 12, 24, 48 and 72 hours following electroporation and in mature adipocytes 11 days after electroporation corresponding to day 6 of differentiation.

**Extended data Figure 3.**
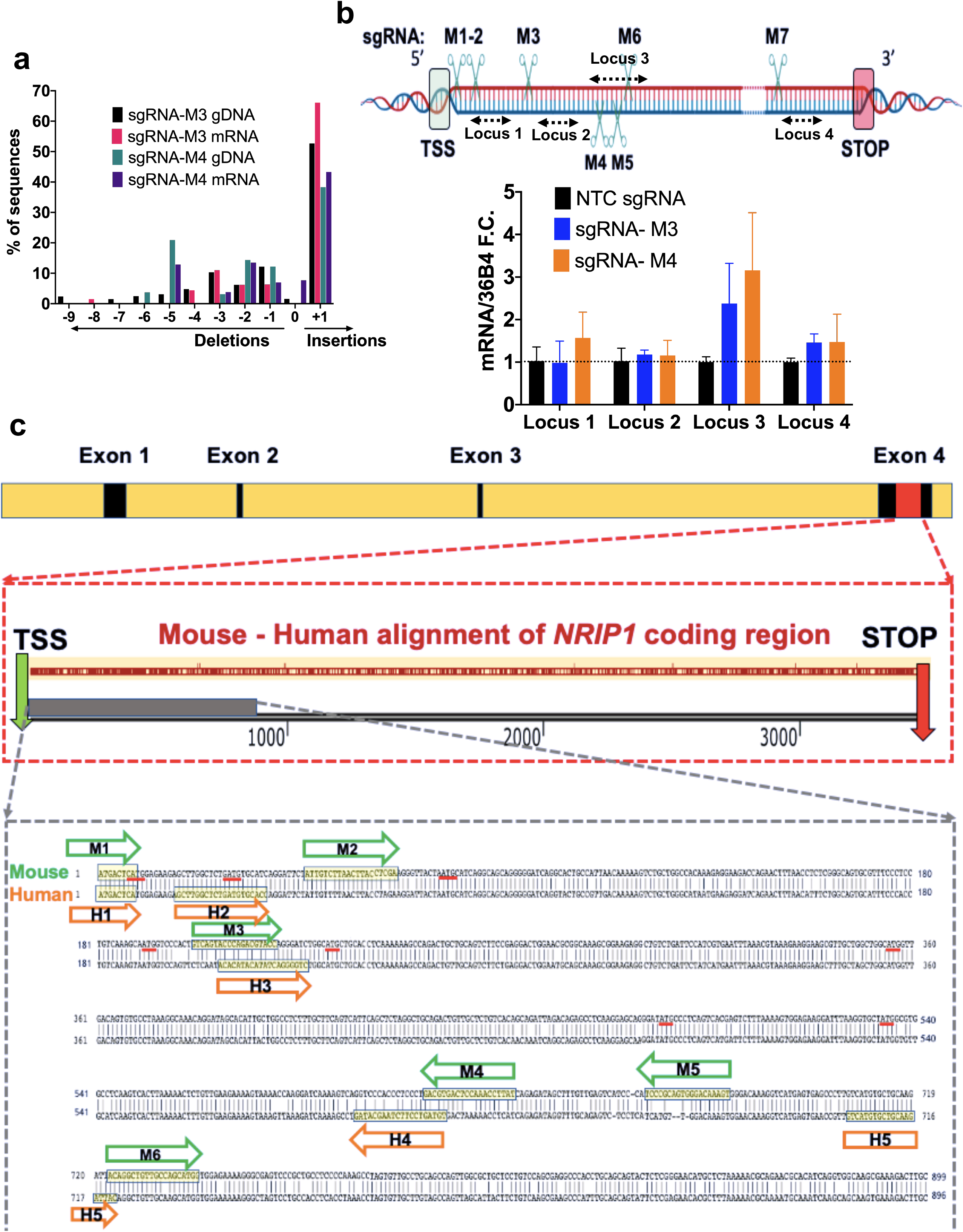
mRNA of *Nrip1* harbors the indels created by the CRISPR-RNPs with little evidence of truncation or degradation of modified *Nrip1* mRNAs. **a.** Comparison of the indel distribution of the genomic DNA and the cDNA after double rDNAse treatment in the template RNA of cells transfected with sgRNA-M3 and sgRNA-M4. **b.** RT-PCR results of the expression of different loci across *Nrip1* cDNA as shown in the map. **c.** Top: *Nrip1* gene includes 4 exons (black) and the coding region (red) is contained in exon 4. Middle: Alignment of the mouse and human coding regions of *NRIP1* that spans 3486bp and 3477bp respectively between the TSS and STOP codons, highlighting the site targeted with sgRNAs M1-6 and H1-5 (gray). Bottom: 5’→ 3’ DNA sequence alignment of the mouse (top strand) and human (bottom strand) of the N-terminus area targeted with the sgRNAs M1-6 for the mouse (green arrows) and H1-5 for the human (orange arrows). Underlined (red) are the ATG sequences in frame or out of frame between the TSS and the first sgRNA that depletes Nrip1 which could potentially serve as alternative translation initiation sites. In panel b error bars denote mean ± standard deviations. n ≥ 3 biological replicates.

**Extended data Fig.4.**
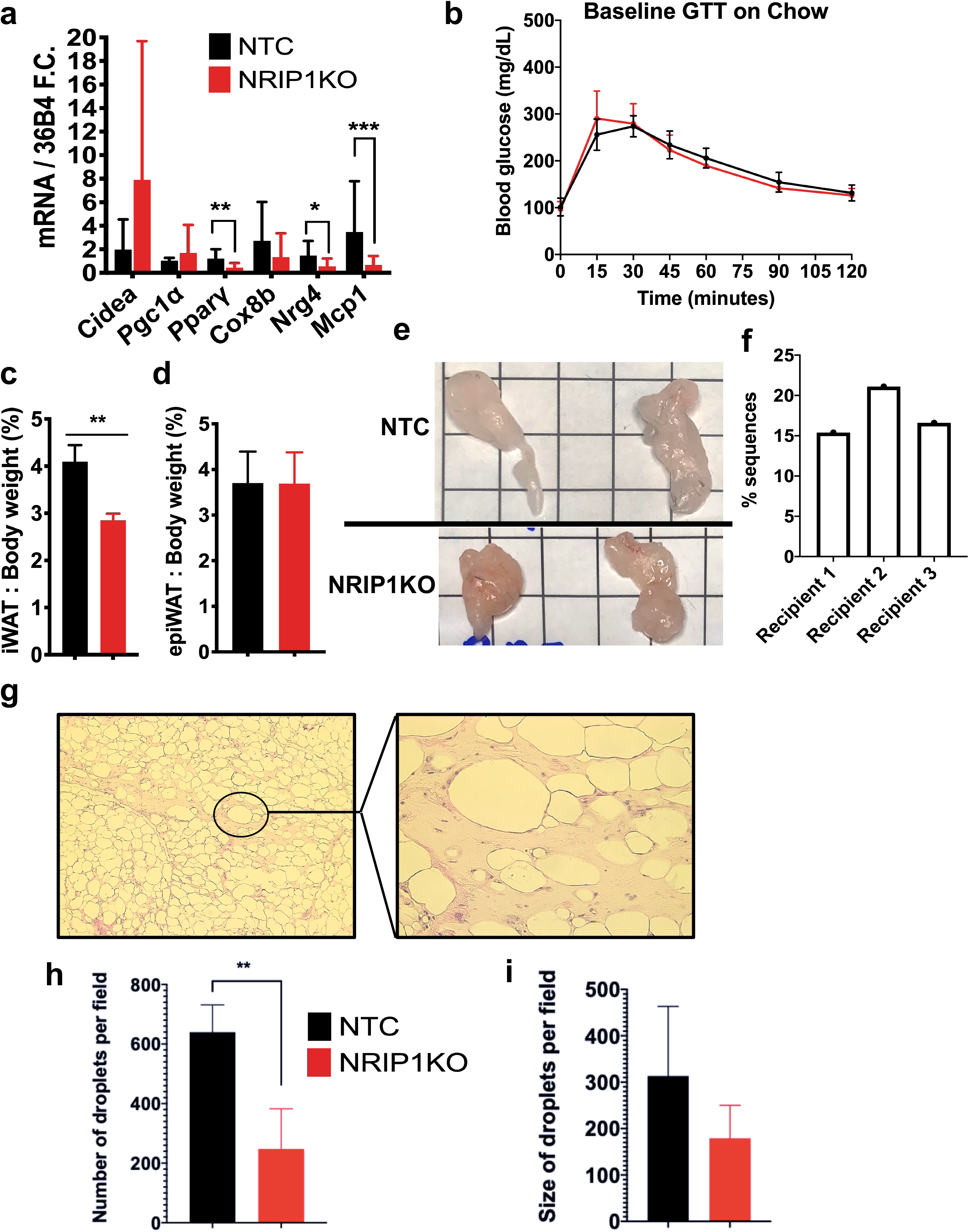
Characterization of mice implanted with Cas9/NTC sgRNA-versus Cas9/sgRNA-M6-treated adipocytes. **a.** RT-PCR results for expression of genes involved in thermogenesis, mitochondrial electron transport chain, lipid and glucose metabolism and the neurotrophic factor Nrg4 prior to implantation. **b.** Baseline GTT after 16-hour fasting in the chow-fed recipients before the implantation of adipocytes. **c**. iWAT over whole body weight percentage. **d**. epiWAT over whole body weight percentage. **e.** Macroscopic images of the whole implants of the recipients after dissection (square = 1cm^2^). **f.** Editing evaluation of homogenized implant sample for each of the NRIP1KO adipocyte recipients. **g.** Histology images stained with H&E of the implant in 5X magnification (left) and 20X magnification (right). **h.** Quantification of the number and **i.** size of lipid droplets in the histology of the livers of the mouse NRIP1KO adipocytes recipients. Panel a: Black = NTC cell implant recipients (n=7); red = NRIP1KO cell implant recipients (n=8), Panel b-h: Black = NTC cell implant recipients (n=4); red = NRIP1KO cell implant recipients (n=3), Bars denote mean, error bars denote mean ± standard deviation. * p < 0.05, ** p < 0.01 by unpaired two-tailed T-test.

## Methods

### Animals and Diets

All animal work was approved by the University of Massachusetts Medical School Institutional Animal Care Use Committee with adherence to the laws of the United States and regulations of the Department of Agriculture. Mice were housed at 20-22 °C on a 12-hour light/12-hour dark cycle with ad libitum access to food and water. C57BL/6J male mice were purchased from Jackson Laboratory for implant studies. C57BL/6J (Jackson Laboratory) male mice were bred for primary preadipocyte cultures. Briefly, 10-week old male mice arrived and were allowed to acclimate for a week prior to any procedures. Mice were implanted with edited primary mouse adipocytes at 11 weeks of age by anesthetizing prior to the implantation procedure using an anesthesia vaporizer chamber with a continuous flow 500 cc/minute of 0_2_ with isoflurane 3% for induction and 1.5% for maintenance. After the cell injections, animals are allowed to wake up and were placed back in clean cages. Mice were maintained on a chow diet for the first 6 weeks, followed by a 60 kcal% high fat diet (Research Diets, D12492i) for the remainder of the experiment from 6-16 weeks post implant. Glucose tolerance tests were performed after 16-hour overnight fasting with intraperitoneal injection of 1g/kg D(+) glucose. Insulin tolerance tests were performed with 0.75IU/kg after 6-hour daytime fasting. Male NOD.Cg-*Prkdc*^*scid*^ *II2rg*^*tm1Wjl*^/SzJ (denoted as NSG) mice were kindly donated by Taconic Biosciences, Inc. NSG mice were implanted with edited primary human adipocytes at 11 weeks of age. Mice were maintained on a chow diet for the first 10 weeks, followed by placing them at thermoneutral with a 60 kcal% high fat diet (Research Diets, D12492i) for the remainder of the experiment from 10 to 15 weeks post implant. Housing under thermoneutrality was achieved by placing the NSG mice at 30°C on a 12-hour light/12-hour dark cycle. Glucose tolerance tests with NSG mice were performed after a 16-hour fast with intraperitoneal injection 2 g/kg D(+) glucose. Whole blood was drawn and placed in EDTA-containing tubes from living mice with submandibular vein punctures under anesthesia as described above and in the end of the study with cardiac puncture. Plasma was extracted with centrifugation of whole blood for 15 minutes at 300 rcf at 4 °C.

### Human Subjects

Abdominal subcutaneous adipose tissue was obtained from discarded tissue following panniculectomy. All subjects consented to the use of tissue and all procedures were approved by the University of Massachusetts Institutional Review Board.

### Primary Mouse Preadipocyte Isolation, Culture and Differentiation to Primary Adipocytes

2 to 3 week old C57BL/6J male mice were euthanized and inguinal fat tissue was harvested (including lymph node) and placed in HBSS buffer (Gibco #14025) plus 3% (w/v) bovine serum albumin (BSA) (American Bioanalytical). The protocol was carried out as described previously^34^ with the following modifications; cells were incubated in 2 mg/mL collagenase (Sigma #C6885) in HBSS BSA 3% (w/v) for 20 minutes to digest the tissue. Cells were cultured to sub-confluence in complete media containing DMEM/F12 media (Gibco #11330), 1% (v/v) Penicillin/streptomycin, 10% (v/v) Fetal bovine serum (FBS) (Atlanta Biologicals #S11550), 100 μg/mL Normocin (Invivogen #Ant-nr-1) at which time they were transfected with RNPs by electroporation and re-plated. Αdipocyte differentiation was induced in the edited cells 24 hours post confluence previously described^34^. Cells grown post differentiation induction were cultured in complete media.

### Primary Human Preadipocyte Isolation, Culture and Differentiation to Primary Adipocytes

Explants from human abdominal subcutaneous adipose tissue from individuals undergoing panniculectomy surgery were embedded in Matrigel and cultured in as previously described^33,42^. Human adipocyte progenitors were transfected with RNPs by electroporation and plated at a density greater than 70% confluence to allow for expansion. Cells were grown to confluence then adipogenic differentiation media was added to induce adipogenesis^33,42^. On day 10 post differentiation, cells were harvested for implantation in NSG mice by treating with 0.5 mg/mL collagenase in 1x Trypsin to detach from culture plates.

### Transfection of Primary Preadipocytes (Mouse and Human) with RNPs

For ribonucleoprotein (RNP) transfection, we used the Neon Transfection System 100 μL Kit (ThermoFisher, #MPK10096) and we prepared a mix consisting unless otherwise specified of sgRNA 40 pmol (Synthego or IDT DNA) purified *Spy*Cas9 protein 30 pmol (PNA Bio, #CP02 or 3xNLS-SpCas9^43^(prepared by the Scot Wolfe laboratory) in Buffer R provided in the Neon Transfection System Kit. The cells were resuspended in Resuspension Buffer R for a final number of 0.5-6 ×10^6^ cells per electroporation. For delivering the RNP complex into primary pre-adipocytes the electroporation parameters used were voltage 1350 V, width of pulse 30 ms; number of pulses 1 unless otherwise specified. The electroporated cells were placed in complete media immediately following transfection, expanded, grown to confluence and differentiated into mature adipocytes for downstream applications. We found these methods improved the viability of preadipocytes and adipocytes and their ability to differentiate over methods reported while our manuscript was in preparation^44^.

### Implantation of Primary Mouse and Human Adipocytes

Primary mouse and human mature adipocytes on day 6 and 10 post differentiation respectively were washed twice with 1xPBS. 0.5 mg/mL collagenase in 1 x trypsin was used to dissociate the cells from the plate. The detached cells are pelleted at 300 rcf for 10 minutes at room temperature. The cells were washed with 1xPBS, pelleted, and the PBS was removed. Cell pellets were kept on ice for a brief time until implantation. Each mouse adipocyte pellet deriving from 1 × 150 mm fully confluent plate was mixed with matrigel (Corning® Matrigel® Growth Factor Reduced Basement Membrane Matrix, Phenol Red-free, LDEV-free # 356231) up to a total volume of 500 μL on ice and the cell and matrigel suspension (500 ± 20 μL) was drawn into a 1 mL tuberculin syringe without the needle. The cell and Matrigel mixture was injected into the anesthetized mouse recipient with a 20 G needle by tenting the subcutaneous subscapular area, inserting the needle into the tented space and injecting at a slow but continuous rate to avoid cell rupture and solidification of the matrigel. The injection site was pinched gently for 1 minute to allow the implant to solidify, followed by withdrawing the needle with a twisting motion. Each C57BL/6J mouse recipient was injected with 2×150mm plates of fully confluent murine adipocytes split into two bilateral injections in the subscapular area. Each NSG mouse recipient received 1 × 150 mm plate split into two 500 μL bilateral subcutaneous injections in the dorsal area as described above.

### DNA Harvest from cells and tissue

At two distinct time-points, 72 hours following transfection and after primary adipocyte differentiation between day 6-10 post differentiation, genomic DNA was isolated from the transfected cells using DNA QuickExtract™ Buffer (Lucigen) in adherence to the manufacturer’s instructions.

### Indel analysis by TIDE and ICE

Genomic DNA was PCR amplified for downstream analysis using locus specific primers designed with MacVector 17.0 and purchased from IDT DNA and Genewiz, spanning the region 800 bp around the expected DSB. For the PCR, Kappa 2x Hot start HiFi mix was used and PCR products were purified using the QIAgen DNA purification kit following the manufacturer’s instructions, and submitted to Genewiz for Sanger Sequencing. Sanger sequencing trace data were analyzed with TIDE and ICE webtools (http://shinyapps.datacurators.nl/tide/, https://ice.synthego.com/#/) that decipher the composition of indels created at the sites of DSBs^45,46^.

### RNA Isolation

Transfected cells were harvested for RNA between day 6-10, post-differentiation depending on the experiment by removing media and washing once with 1xPBS, and adding Trizol reagent to lyse the cells. The protocol for RNA isolation was performed according to manufacturer’s instruction with the following modifications; 1μl of Glycol blue (Invitrogen #AM9516) was added to the isopropanol to precipitate the RNA and was either stored overnight at −20°C or placed on dry ice for 2 hours. The isolated RNA was resuspended in RNase free water, then treated with recombinant DNaseI (DNA-free DNA removal kit, Ambion) according to the manufacturer’s instructions. RNA concentrations were determined by Nanodrop 2000.

### RNA Isolation of Pulverized Tissue/Tissue Piece

Tissue was isolated from the mice and frozen in liquid N2. For RNA isolation, tissue was pulverized in liquid N2, or a piece approx. 100 mg in size was put in a 2 mL tube with screw cap and 1mL of Trizol. Tissue was placed in the Qiagen TissueLyser and homogenized for 3 cycles of 3 minutes at 30Hz. The Trizol and tissue lysate were placed in a new tube, and centrifuged for 10 minutes at 4°C to separate any lipid from the homogenate. Once the homogenate is separated from the lipid, the remaining isolation is carried out according to manufacturer’s instructions.

### RT-PCR

0.5-1 μg of RNA was used in 20 μL reaction with Bio-Rad iScript cDNA kit according to manufacturer’s protocol to synthesize cDNA. cDNA was diluted by adding 80 μL of water to the reaction and 5 μL of cDNA template was used for RT-PCR with Bio-Rad Sybr Green Mix and gene specific primers for a final concentration of 0.3 μM primers. Expression of genes was determined by comparing gene expression levels of target gene compared to housekeeping gene 36B4 and RPL4 for murine and human samples respectively. mRNA expression was analyzed with the ΔΔCT method.

### Protein Isolation

Cells grown in culture dishes were washed once with 1 x PBS at room temperature, followed by adding boiling 2% SDS (w/v) with 1 X HALT protease inhibitors and scraping to lyse the cells. Tissue pieces were prepared for western blots by homogenizing a piece approximately 100 mg in Radioimmunoprecipitation Assay (RIPA) buffer with 1x HALT protease inhibitors in the Qiagen TissueLyser and homogenized for 3 cycles of 3 minutes at 30Hz. Tissue and cell lysates prepared with 2% SDS (w/v) buffer or RIPA buffer were sonicated at 60% amplitude with a probe sonicator tip for 30 seconds at room temperature. In figure S2, mouse cells were lysed as described above at different time-points after transfection and for timepoint 0 hours, after the electroporation the transfection mix consisting of cells and RNPs in Buffer R was centrifugated at 300 rcf. The cell pellet was lysed as described above while the supernatant (sup) was also collected for use as positive control (30pmols of SpyCas9) in the western blot. Protein concentration determination of the tissue and cell lysates was performed using a bicinchoninic acid kit (BCA Protein Assay Kit, Pierce). Cell lysates used in immunoprecipitation reaction were prepared in non-denaturing NP-40 buffer (20mM Tris HCl pH 8.0, 137mM NaCl, 1% (v/v) Nonident P-40 (NP_40), 2mM EDT) containing 1X HALT protease inhibitors by washing once with 1 x PBS, adding NP-40 buffer and scraping, followed by a 4 °C incubation for 30 minutes to 1 hour with gentle agitation. Cell lysates were centrifuged at 4 °C for 10 minutes at 16,100 rcf and the infranate was collected and used. Protein concentrations were determined on the lysates using Pierce BCA Kit. Protein samples were prepared for running on 7.5-12 % SDS-PAGE mini gels at a final concentration of 1mg/mL protein, 1x Laemmli loading buffer (BioRad) with 2.5 %(v/v) β-Mercaptoethanol, followed by placing in a heat block at 95 °C for 10 minutes.

### Triglyceride Assay

For the liver triglyceride assay, we used the Triglyceride Colorimetric assay kit (Cayman Chemical, #10010303). The lysate was prepared by mixing 50 mg of pulverized whole liver with 1.5 mL of the NP-40 lysis buffer and homogenized in the Qiagen TissueLyser with 3 cycles 3 minutes at 30 Hz. The assay ran according to manufacturer instructions with a sample dilution of 1:5.

### Human Adiponectin

Human adiponectin was measured in the plasma of NSG mice was measured using a human-specific adiponectin ELISA from Invitrogen (KHP0041).

### Histology

Approximately 0.5 cm^2^ of the implant tissue and two 0.5 cm^2^ liver pieces from two different lobes per recipient were randomly selected and fixed, followed by processing at the UMass Medical School Morphology Core. Photos of the tissues were taken with an LEICA DM 2500 LED inverted microscope at indicated magnification. Fiji/ImageJ was used to quantify lipid content in H&E images. 4 images per section, 2 sections per liver, were projected into a single montage. The montage was converted from RGB to 8 bit, contrast enhanced, thresholded and binarized. The processed montage was reconverted into individual images and lipid droplets quantified for each image using the particle analysis function (number, size, % of area covered).

### Western Blotting and Immunoprecipitation

Protein lysates were run on 7.5 and 12% SDS-PAGE or Mini-Protean TGX stain-free pre-cast protein gels, followed by transferring the proteins to Nitrocellulose. Unless otherwise stated, a total of 20 μg of protein lysate was loaded per well. Nitrocellulose membranes were blocked using 5% (w/v) Non-fat milk in Tris buffered saline with 0.1% (v/v) Tween-20 (TBST) for 1 hour at room temperature. Primary antibody incubations were carried out in 5% (w/v) BSA in TBST at the following antibody concentrations: UCP1-Abcam#10983, 1:700; Rip140-Millipore #MABS1917, 1:1000, Tubulin-Sigma #T5168, 1:4000; GAPDH-Cell Signaling #21185, 1:1000; SpyCas9-Cell Signaling #19526S 1:5000. Blots and primary antibodies were incubated overnight with a roller mixer at 4°C.

Membranes were washed with TBST prior to secondary antibody incubations. HRP-conjugated secondary antibodies were diluted with 5% BSA (w/v) in TBST at 1:5,000-10,000 for 45 minutes at room temperature with constant shaking. Membranes were washed in TBST, followed by incubating with Perkin Elmer Western Lightning Enhance ECL. The Bio-Rad Chemi-Doc XRS was used to image the chemiluminescence and quantifications were performed using the system software, or Image J. Immunoprecipitation was performed with NP-40/Halt protein lysates. Briefly, 250 μg of protein lysates were pre-cleared using 50:50 Protein-A Sepharose/NP-40 buffer/1x HALT protease inhibitors for 2 hours at 4°C with end over end mixing. After 2 hours, the lysate/Protein A Sepharose was centrifuged for 5 seconds to pellet the Sepharose, and the lysate was transferred to a new tube. 5 μg of Antibody (Rabbit Non-Immune IgG, Millipore #12-370, or Rabbit anti-Nrip1, Abcam #Ab42126) was added to the lysates and they were incubated overnight at 4 °C with end over end mixing. Antibody/antigen was pulled down by adding 50:50 Protein A Sepharose/NP-40 buffer/1x HALT protease inhibitors for an additional 2 hours at 4°C with end over end mixing. Protein/Antibody/Protein A Sepharose complexes were washed by centrifuging briefly, removing the supernatant and washing the pellet with NP-40 buffer containing protease inhibitors. The captured proteins were eluted from the Sepharose by adding 40 μL of 1xLaemmli buffer containing 2.5% (v/v) β-Mercaptoethanol, vortexing the Sepharose mixture, followed by boiling at 95°C for 10 minutes. All eluted proteins were run on the gel, transferred to nitrocellulose, and immunoblotted as described above.

### Plasmid construction

The pCS2-Dest plasmid with CMV promoter expressing SpyCas9 (Addgene # 69220), and sgRNA expressing plasmid (Addgene #52628) were a gift from Dr. Scot Wolfe lab. To clone NRIP1 targeting and non-targeting sgRNAs, oligo spacers with BfuAI overhangs (purchased from IDT) were annealed and cloned into the BfuAI-digested sgRNA plasmid. Lonza pmaxGFP LOT 2-00096 was used to test transfection of these plasmids in various concentrations to determine the efficient dosage range (0.5–1.5μg) and the electroporation conditions (1350 V, 30ms, 1 pulse) for the delivery and GFP expression was evaluated with EVOS FL fluorescent microscope (Thermo Fisher Scientific).

### Statistical analysis

All comparisons are between two groups and student unpaired two-tailed T-Test was performed for the *p* values. In data that did not follow Gaussian distribution, standardization preceded the statistical analysis. * *p* < 0.05, ** p < 0.01, *** p < 0.001.

### Additional online tools and Software

For the mapping of exons on the Nrip1 gene we used IGV_2.5.3. For the Design of sgRNAs we used a combination of the Broad Institute sgRNA designer, CHOPCHOP and the online sgRNA checkers by Synthego and IDT. For the design of genomic DNA primers we used MacVector 17.0. For the alignment of the Sanger Sequencing traces and the human and mouse coding region we used SnapGene Viewer 5.1.6 and NCBI nucleotide blast. For the design of RT-PCR primers we used Primer Bank (https://pga.mgh.harvard.edu/primerbank/). For the prediction of off-target editing sites, we used Standard nucleotide blast by NCBI to determine the first three potential binding sites with a PAM sequence downstream. For the data graphing, we used Prism GraphPad 8.

**Table 1.**
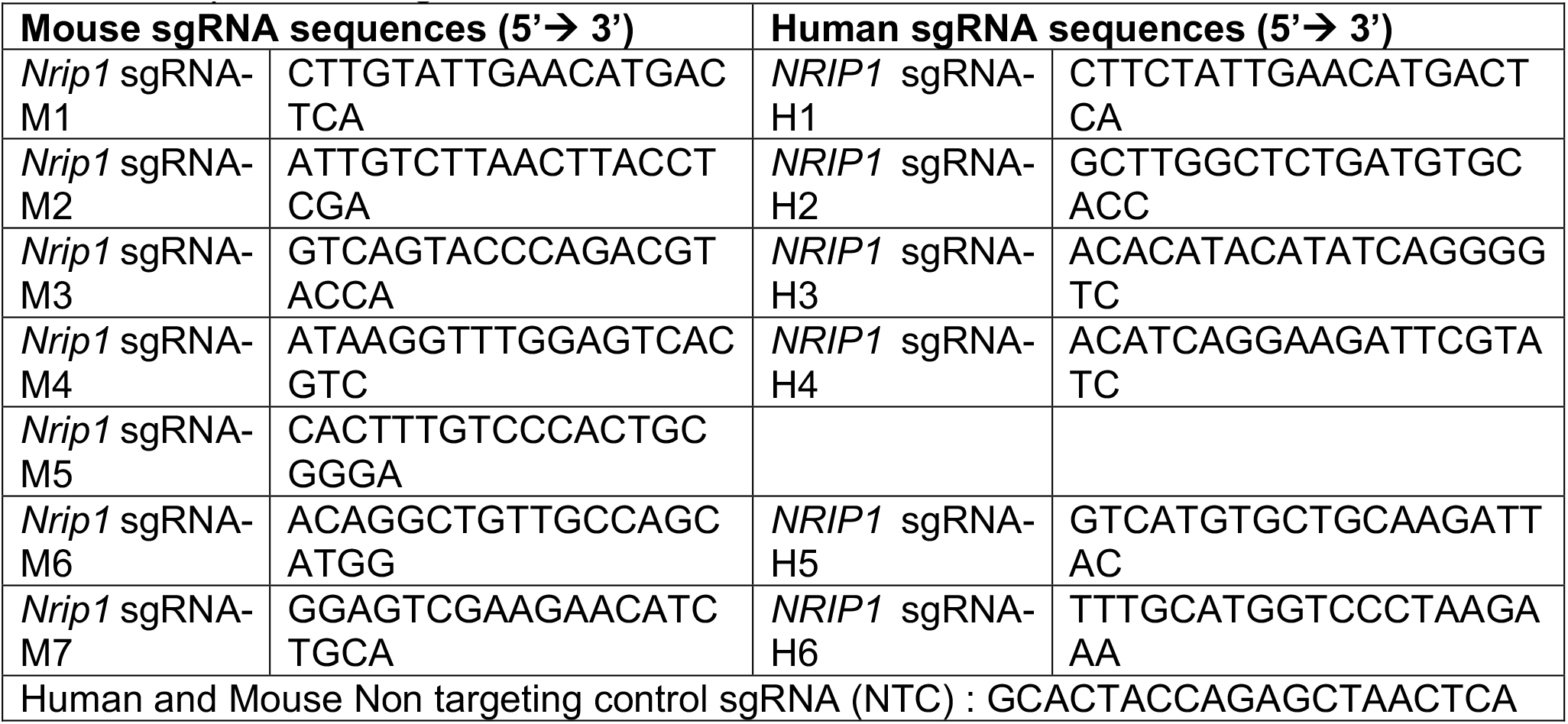
Sequences of sgRNAs.

**Table 2.**
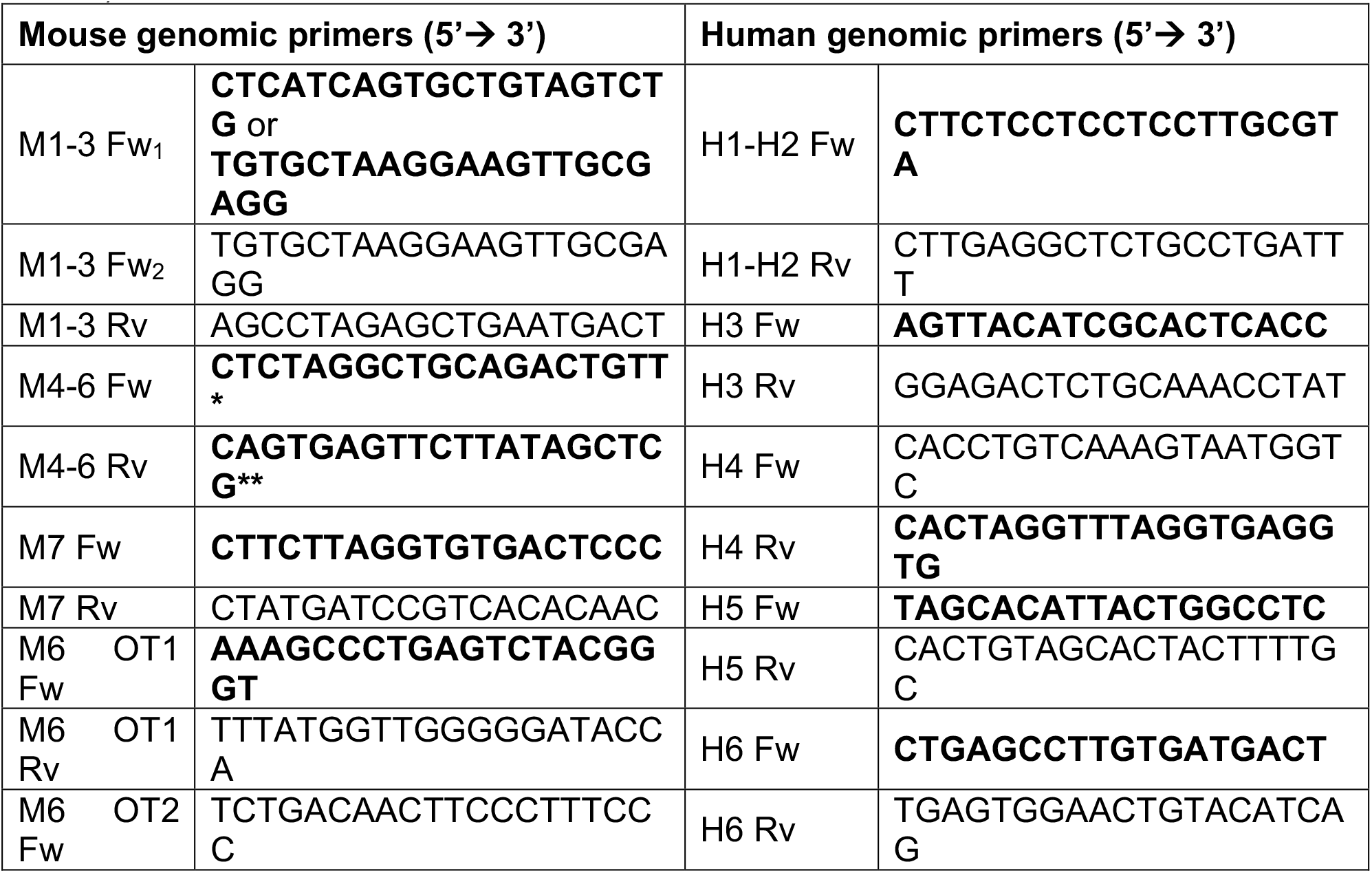
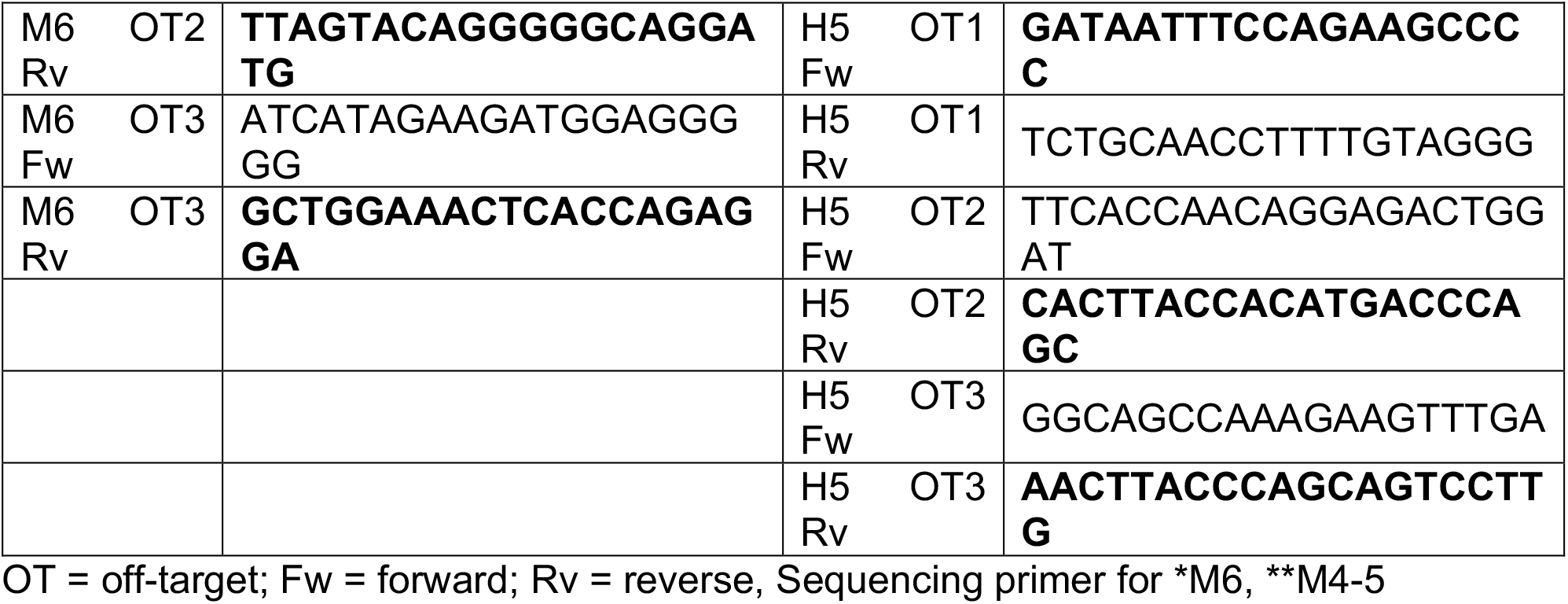
Primers for genomic DNA PCR and Sanger sequencing (sequencing primers bolded)

**Table 3.**
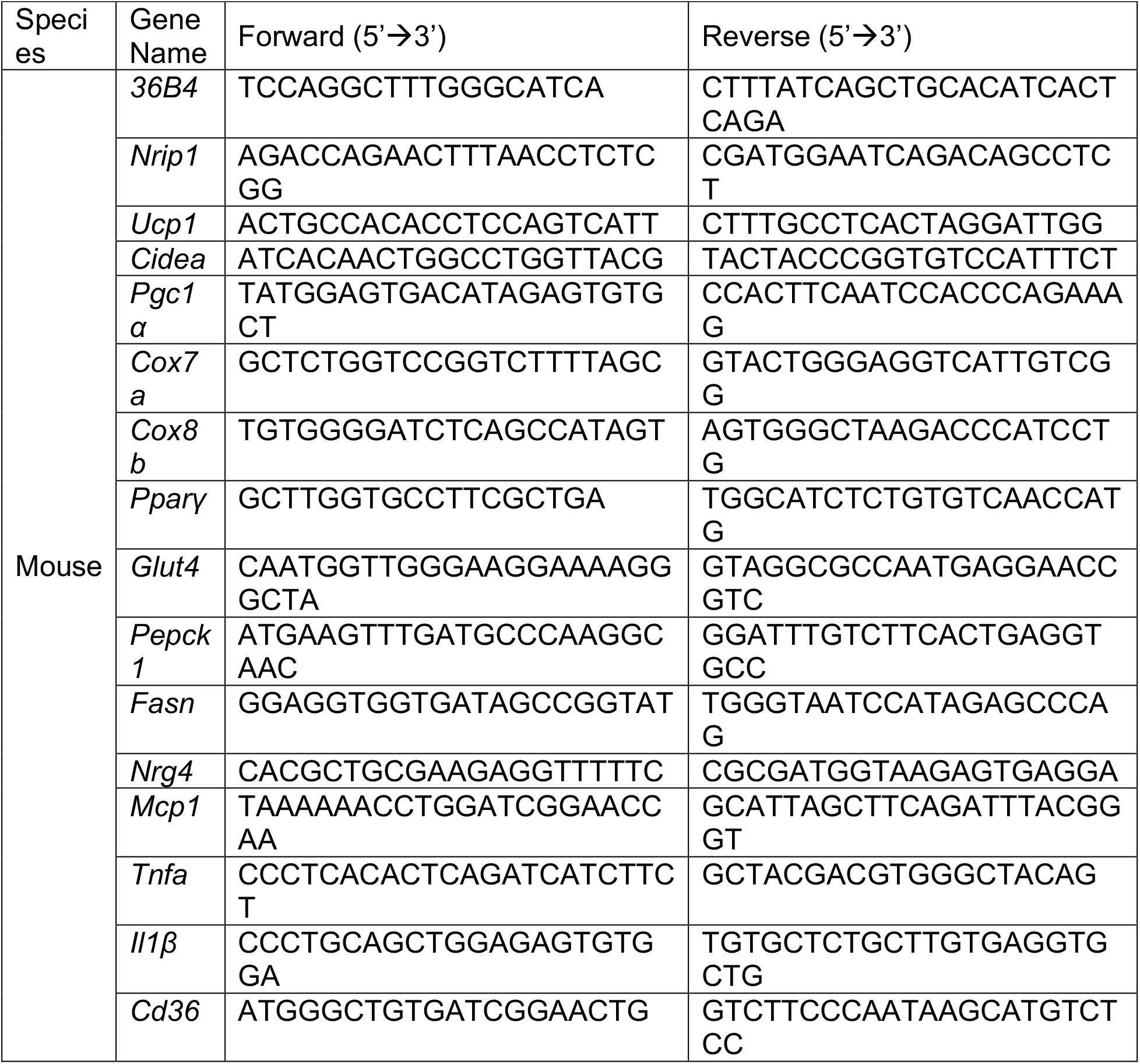
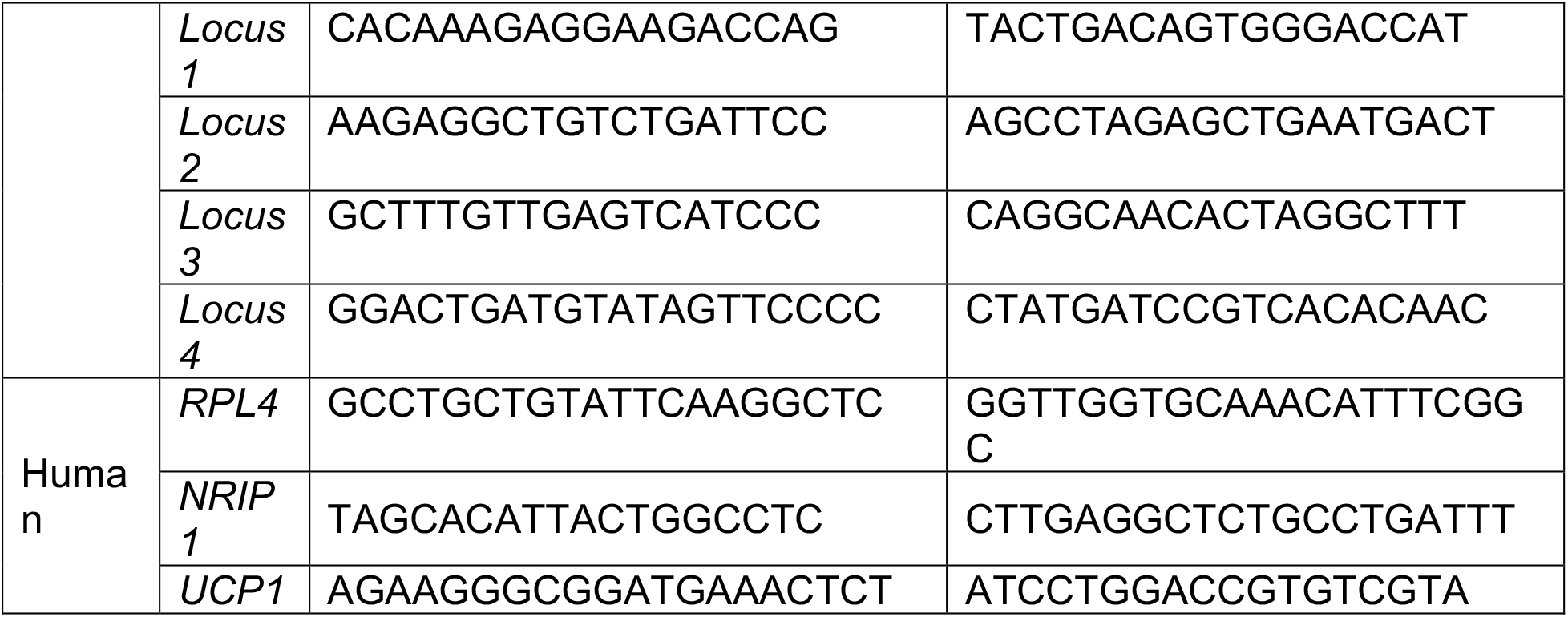
RT-PCR Primers.

## Notes

### Competing Interest Statement

The authors have declared no competing interest.

